# Does Vibrotactile Stimulation of the Auricular Vagus Nerve Enhance Working Memory? A Behavioral and Physiological Investigation

**DOI:** 10.1101/2024.03.24.586365

**Authors:** Gansheng Tan, Josh Adams, Kara Donovan, Phillip Demarest, Jon T. Willie, Peter Brunner, Jenna L. Gorlewicz, Eric C. Leuthardt

## Abstract

**Background:** Working memory is essential to a wide range of cognitive functions and activities. Transcutaneous auricular VNS (taVNS) is a promising method to improve working memory performance. However, the feasibility and scalability of electrical stimulation are constrained by several limitations, such as auricular discomfort and inconsistent electrical contact.

**Objective:** We aimed to develop a novel and practical method, vibrotactile taVNS, to improve working memory. Further, we investigated its effects on arousal, measured by skin conductance and pupil diameter.

**Method:** This study included 20 healthy participants. Behavioral response, skin conductance, and eye tracking data were concurrently recorded while the participants performed N-back tasks under three conditions: vibrotactile taVNS delivered to the cymba concha, earlobe (sham control), and no stimulation (baseline control).

**Results:** In 4-back tasks, which demand maximal working memory capacity, active vibrotactile taVNS significantly improved the performance metric *d*’ compared to the baseline but not to the sham. Moreover, we found that the reduction rate of *d*’ with increasing task difficulty was significantly smaller during vibrotactile taVNS sessions than in both baseline and sham conditions. Arousal, measured as skin conductance and pupil diameter, declined over the course of the tasks. Vibrotactile taVNS rescued this arousal decline, leading to arousal levels corresponding to optimal working memory levels. Moreover, pupil diameter and skin conductance level were higher during high-cognitive-load tasks when vibrotactile taVNS was delivered to the concha compared to baseline and sham.

**Conclusion:** Our findings suggest that vibrotactile taVNS modulates the arousal pathway and could be a potential intervention for enhancing working memory.

**Highlights:**

- Vibrotactile stimulation of the auricular vagus nerve increases general arousal.
- Vibrotactile stimulation of the auricular vagus nerve mitigates arousal decreases as subjects continuously perform working memory tasks.
- 6 Hz Vibrotactile auricular vagus nerve stimulation is a potential intervention for enhancing working memory performance.

## 1. Introduction

Working memory is a limited-capacity system that manipulates information and translates it into behaviors. It is the foundation of various cognitive functions, such as planning and learning. Critically, working memory deficits are common in numerous neurological disorders, affecting the quality of life for millions^1^. Despite its importance, improving working memory presents a formidable challenge with brain stimulation methods because of its distributed nature^2^.

To address this, we propose modifying a top-down modulatory mechanism, arousal, to improve working memory performance^3^. In this context, the vagus nerve presents an appealing target. The Vagus nerve primarily contains afferent fibers that project to brain areas, including the nucleus tractus solitarius (NTS), locus coeruleus (LC), and basal forebrain structure, which is summarized in ***Figure 1*a**^1, 4^. These sites are involved in regulating arousal and attention and, therefore, critical to working memory^5^. Vagus nerve stimulation (VNS) is shown to increase arousal levels in animal studies and has cognitive benefits in human studies^6, 7, 8, 9, 10, 11, 12, 13, 14^.

In recent years, transcutaneous auricular vagus nerve stimulation (taVNS) has emerged as a noninvasive alternative, garnering attention for its potential in treating a range of clinical conditions. The auricular branch of the vagus nerve runs superficially and has cutaneous receptive fields in the external ear, thereby allowing for noninvasive techniques to stimulate the vagus nerve^4, 13^. Specifically, the cymba concha is thought to be almost entirely innervated by the auricular branch of the vagus nerve, while other areas, such as the tragus and crus of the helix, may engage some vagus fibers^15, 16^. Despite the potential cognitive benefits of taVNS, there are significant barriers to extending its application towards broader clinical and non-clinical uses, including the associated discomfort of electrical stimulation and the technical challenge of achieving consistent electrical contact in the ear.

Vibrotactile stimulation represents a practical and non-aversive approach to activating the vagus nerve. Following an animal study demonstrating the anti-inflammatory effect of mechanical stimulation of the vagus nerve, Addorisio et al. showed that vibrotactile stimulation administered to the cymba concha region of the outer ear elicited VNS-mediated anti-inflammatory effects^17, 18^. Building on this foundation, we hypothesized that vibrotactile stimulation of the auricular vagus nerve would improve working memory through neuromodulation of arousal levels.

## 2. Material and methods

### 2.1 Subjects

The study enrolled 22 healthy individuals between 18 and 45 years of age without a significant past medical history and with no known neurological deficits. Participants were required to abstain from alcohol and caffeine consumption starting one day before the study. The study was performed in accordance with the Declaration of Helsinki, the International Conference on Harmonization Good Clinical Practice Guidelines, and the applicable United States Code of Federal Regulations. The study was approved by The Washington University in St. Louis Institutional Review Board (IRB ID#: 202101161). Written informed consent was obtained from all study participants. Out of the 22 participants initially recruited, two were excluded from the analysis (see supplementary Figure 1 for details).

### 2.2 Intervention

The vibrotactile taVNS system comprises parts fabricated with stereolithographic additive manufacturing (***Figure 1*b-c**). The system includes an ear cup (b1), a cushioning base (b2) that provides soft contact with the head, a T-head bolt (b5) that connects the contact tip (b3) and the ear cup (b1) via ear cup covers (b4). An eccentric rotating mass (ERM) vibration motor of 5mm diameter (Vybronics Inc, Brooklyn, NY) is held in at the contact tip (b3). The motor’s power cable passes through the T-head bolt (b5). The T-head bolt allows individual adjustment of the depth of the contact. In addition, rotating the T-head bolt allows switching the stimulation target between the concha area and the earlobe. A digital stimulation box (g.STIMBOX, g.tec medical engineering, Austria) supplied 5V power at 6 Hz to the eccentric Rotating Mass vibration motor. To the best of our knowledge, our study is the first investigation of the effects of vibrotactile taVNS on working memory. Lacking prior research on the impact of different stimulation frequencies of vibrotactile taVNS, we selected a frequency of 6 Hz for this pilot study. This decision was informed by literature indicating that theta oscillation, within the 4-7 Hz range, may play a crucial role in supporting working memory functions^19, 20^. An elastic strap stabilizes the vibrotactile device on the head (**Figure *1*c**). Participants generally found the vibrotactile device comfortable and the vibration sound tolerable (***Figure 1*d**).

**Figure 1.**
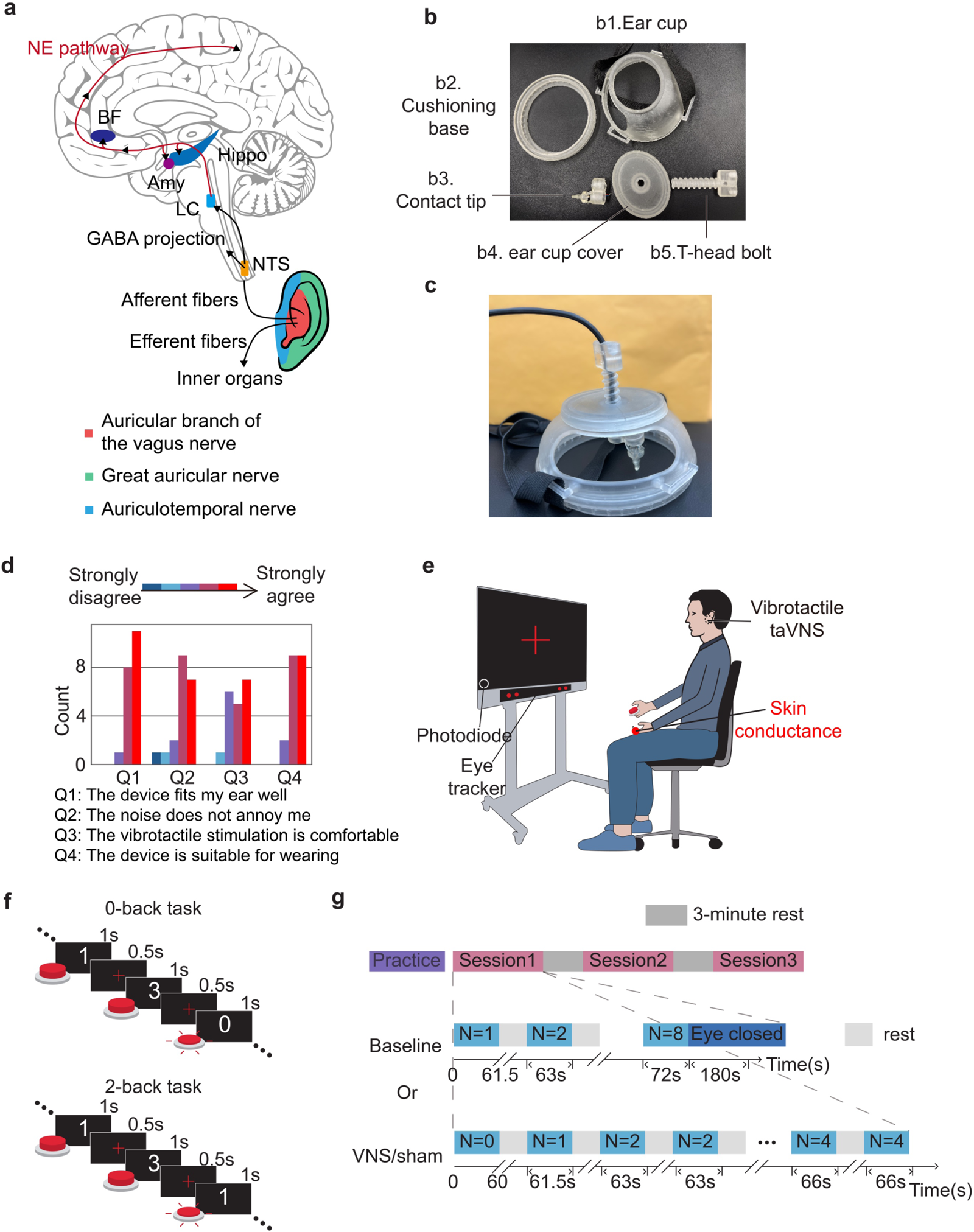
Vibrotactile taVNS system and study design. **a**. Illustration of afferent vagal pathway^1, 4^. The colored area in the ear represents the parts of the ear that are innervated by the auricular branch of the vagus nerve, the great auricular nerve, and the auriculotemporal nerve. **b**. 3D-printing parts of the vibrotactile device. **c**. The assembly of the vibration device. An RCA cable goes through **b5** and supplies the vibration motor. **d**. Survey results about the experience of wearing the vibrotactile stimulation device. **e**. Experiment platform. The participant sits on a chair and holds a button with the dominant hand. **f**. Illustration of N-back task used in this study. For the 0-back task, participants were asked to respond with a button press when they saw 0. In the 2-back task, participants were instructed to respond when they thought the currently presented digit was the same as the digit presented two trials ago. **g**. N ranges from 1 to 8 in the baseline session. The VNS and sham sessions included eight blocks, each with one block for the 0-back task, one for the 1-back task, and two for the 2, 3, and 4-back task.

### 2.3 Study design

This study used N-back tasks to assess working memory. Subjects responded with a button press when they thought the currently presented digit was the same as the digit presented N trials prior (***Figure 1*f**). The digits presented in the N-back task were randomly selected from 4 digits: 0, 1, 2, and 3. Participants completed three sessions of N-back tasks: a baseline session, a sham session, and a VNS session, with the order of sessions randomized (***Figure 1*g**). To mitigate potential confounding factors related to fatigue or residual effects of stimulation, participants rested for 3 minutes between sessions. This protocol was informed by findings from an animal study, which suggest that both cortical and behavioral activations elicited by VNS returned to baseline levels within 30 s^21^. During the baseline session, the participant completed 8 blocks of N-back tasks with N ranging from 1 to 8. Between two blocks, subjects were given a mandatory minimum rest period of 10 seconds, after which they could proceed to the next block by pressing a button. Each block includes 40 probes, that is, digit presentations, when the subject needs to decide whether to press the button. Before the first probe, N digits were presented to load the participant’s working memory. Throughout the task, digits were displayed in white against a black screen background for 1 second each (***Figure 1*f**).

Between the presentation of two digits, a red fixation cross was displayed in red for 0.5 seconds. During the VNS or sham session, the participant completed 8 blocks of N-back tasks with N ranging from 0 to 4. The 0-back task in VNS or sham session was designed as a control for attention. At the end of the baseline session, subjects were asked to rest and close their eyes. Data collected from participants while they closed their eyes was used to ensure eye-tracking and skin conductance recording validity.

During the VNS session, vibrotactile stimulation was delivered to the cymba concha area of the outer ear. In concordance with previous studies, we selected the earlobe as the sham stimulation target^22^. In VNS and sham sessions, subjects were stimulated when performing the N-back tasks. The timing of the stimulation was synchronized with digit presentations: it was activated while the digit was displayed and deactivated during the presentation of the fixation cross. This protocol was designed to reduce the adaptation of the cutaneous mechanoreceptors. During the baseline session, subjects wore the vibrotactile taVNS device that contacted the earlobe. Before the study, the participants received instruction from the experimenter and practiced the N-back task until they were familiar with it.

### 2.4 Outcome measures

We recorded behavioral responses using a push-button sensed by a multimodal trigger box (g.TRIGBOX, g.tec medical engineering, Austria), galvanic skin conductance using a g.USBAMP research amplifier (g.tec medical engineering, Austria), and pupillometry using a Tobii Pro Fusion eye tracker (Tobii, Sweden). We acquired pupil diameter at 120

Hz and a resolution of 0.1 mm. The eye tracker was attached at the bottom of the display monitor. Before the N-back tasks, we adjusted the monitor’s orientation and height to ensure the eye tracker camera captured the eyes. We then used Tobii Eye Tracking software to calibrate the eye tracker. These multimodal data were collected and synchronized with *bci2000*, an open-source software system^23^. Eye tracking data were successfully collected for 17 subjects due to logistical constraints.

#### 2.4.1 Primary outcome measures

We derived hit rate, false alarm rate, reaction time, and *d*’ from the multimodal trigger box to evaluate performance. The primary behavioral outcome measure is d prime (*d*’), representing the sensitivity to distinguish between true positives (hits) and false positives (false alarms). We calculated *d*’ as follows:

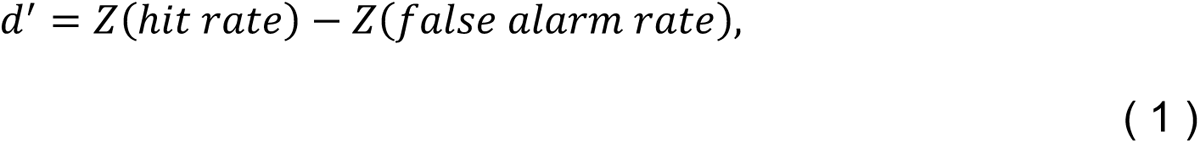

where Z is the inverse of the cumulative Gaussian distribution function. We used skin conductance level and pupil diameter as physiological proxies of arousal.

#### 2.4.2 Event-related skin conductance

We recorded skin conductance at a sampling rate of 1200 Hz and resampled it at 10 Hz for fast processing. We used a 6th-order Butterworth low-pass filter with a cut-off frequency of 2 Hz to remove high-frequency artifacts in skin conductance data. For event-related analysis, we rescaled the skin conductance to a 0 to 1 range and aligned it to the task onset for each N-back task block.

#### 2.4.3 Tonic skin conductance analysis

We used a convex optimization approach to decompose the skin conductance signal into tonic and phasic components, which represent, respectively, general arousal and sympathetic activities^24^. To compare across subjects and sessions, we sampled the skin conductance level every 10 seconds and z-scored it based on the value in the 1-back task within each session. To bridge the behavioral effect and physiological effects of vibrotactile taVNS, we calculated the difference in skin conductance level between the sham/VNS session and baseline session for each difficulty. Next, we grouped performance metrics based on the difference.

#### 2.4.4 Pupillometry analysis

We removed invalid pupil measurements that were induced due to blinks. Blink events were detected when the pupil diameter change rate surpassed 0.3 mm/ms. The blink was assumed to last no longer than 500 ms^25^. If an eye blink offset was not found within 500 ms following an onset, we defined it as 500 ms after the last detected eye blink onset. We added a safety margin of 15 ms around the eye blinks to account for potential pre-blink contamination of the pupillometry measurements. Finally, we resampled the remaining valid pupillometry measurement at 30 Hz for subsequent processing.

To compare the temporal dynamics of pupil diameter across conditions (VNS, sham, and baseline), we first normalize the data. This was done by z-scoring to enable the grouping of pupil diameter based on the conditions. The presentation of the digit and fixation cross induced a periodic component with a 1.5 s cycle into the subject’s pupil diameter. We applied a 0.1 Hz low-pass filter on the linearly interpolated pupil diameter to remove this confounding component and extract the general arousal component.

Furthermore, to elucidate the impact of vibrotactile taVNS on arousal across different N-back task difficulties, we first interpolated and filtered the preprocessed pupil diameter using a 0.1 Hz low-pass filter. Subsequently, we corrected the filtered pupil diameter by subtracting the mean of the filtered pupil diameter during the 1-back task for each session.

### 2.5 Statistical analysis

To titrate working memory performance as a function of difficulty, a non-linear four-parameter log-logistic function is used to model *d*′. In equation *(2)*, the parameters *d* and *c* represent the upper and lower bounds, respectively. The parameter *e* represents the effective dose for a 50% response, that is, the difficulty that results in a *d*′ halfway between the upper and lower limits. The parameter *b* represents the steepness of the curve. Wald test is used to test the significance of the four parameters. To find out the maximum working memory span, T-tests are used to determine if d’ is significantly higher than the parameter c.

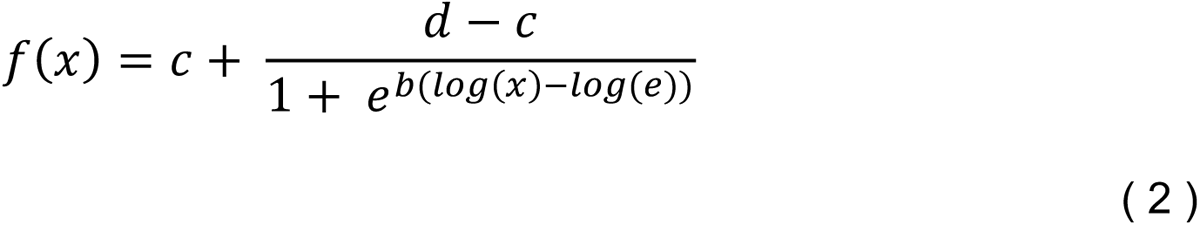

Wald test was used to test the significance of the four parameters. In addition, we used T-tests to see if *d’* at *N* = 3 and *N* = 4 are close to the parameter *c* in equation *(2)*. We used Wilcoxon signed rank tests to compare *d*’ and reaction time between sessions when the maximal working memory span is reached (i.e., N=4). We used Linear mixed-effects models and linear regression models to regress *d*’ and reaction time. Akaike information criterion was used to select the best model. T-tests using Satterthwaite approximation of the degree of freedom were used to identify significant effects. The statistical analyses were performed in R and Python using *lmerTest* package^26^.

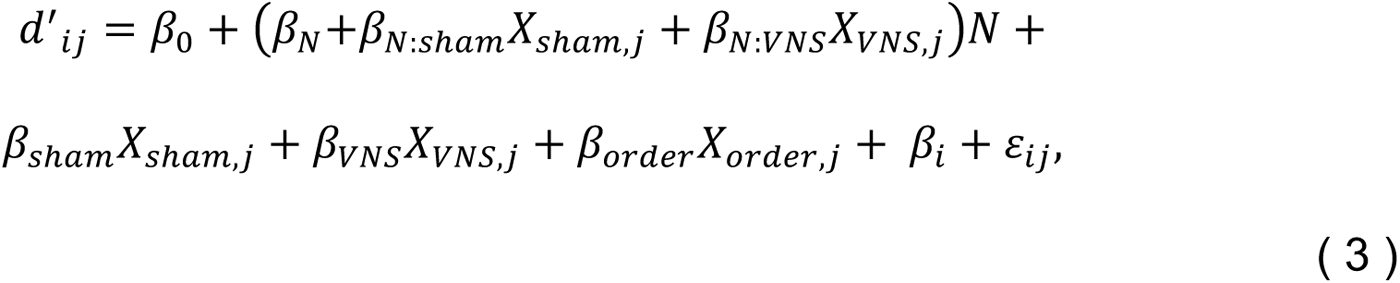

where *i* represents *i^th^* subject and *j^th^* represents *j^th^* session. *X_sham,j_* if the *j^th^* session is sham session, otherwise 0. *X_VNS,j_* session is VNS session, otherwise 0. *N* represents the difficulty, that is, N of N-back. *X_order, j_* represents the order of the *j^th^* session. ε is the normally distributed noise.

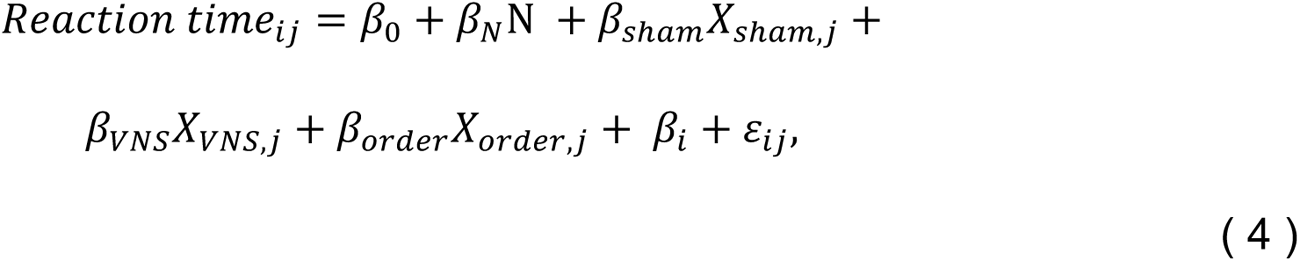

where *i* represents *i^th^* subject, and *j* represents *j^th^* session. *X_sham,j_, X_VNS,j_, X_order,j_*, and ε have the same notation. Scaled residuals are reported to assess the model’s fit, which is residuals divided by an estimate of their standard deviation. To test if the decrease in *d*’ with increasing difficulty in the VNS session was smaller than that in the sham session, we performed a directional post-hoc z-test on *β_N:VNS_* and *β_N-Sham_* using the *multcomp* package in R^27^.

We used permutation tests to compare normalized skin conductance level and normalized pupil diameter between sessions because the p-value is negatively associated with the sample size in T-tests. We calculated observed t-statistics and t-statistics after permutation. The p-value was derived as a proportion of permuted t-statistics greater than observed t-statistics.

## 3. Results

### 3.1 Behavioral outcome at baseline

To identify the dynamic range of task performance, we titrated the N-back task performance with N ranging from 1 to 8 during the baseline session. The hit rate and *d*’ decreased asymptotically as N increased (**Figure *2*a** and **Figure *2*b**). The 4-parameter log-logistic regression model fits well *d*’ (mean residual = 0, residual standard error = 2.5, **Figure *2*c**). The model showed that as difficulty increases, d’ approaches a lower bound representing d’ when the subject presses the button randomly. T-tests showed that *d*’ at n=3 is significantly higher than estimated parameter *c* in the 4-parameter log-logistic regression model (t = 2.71, p-value < 0.01), while *d*’ at n=4 is not (t = 0.94, p-value = 0.18). These results show that subjects actively performed N-back tasks and that the maximum working memory span was most likely reached at 4-back, which aligns with studies of working memory capacity^28^.

**Figure 2.**
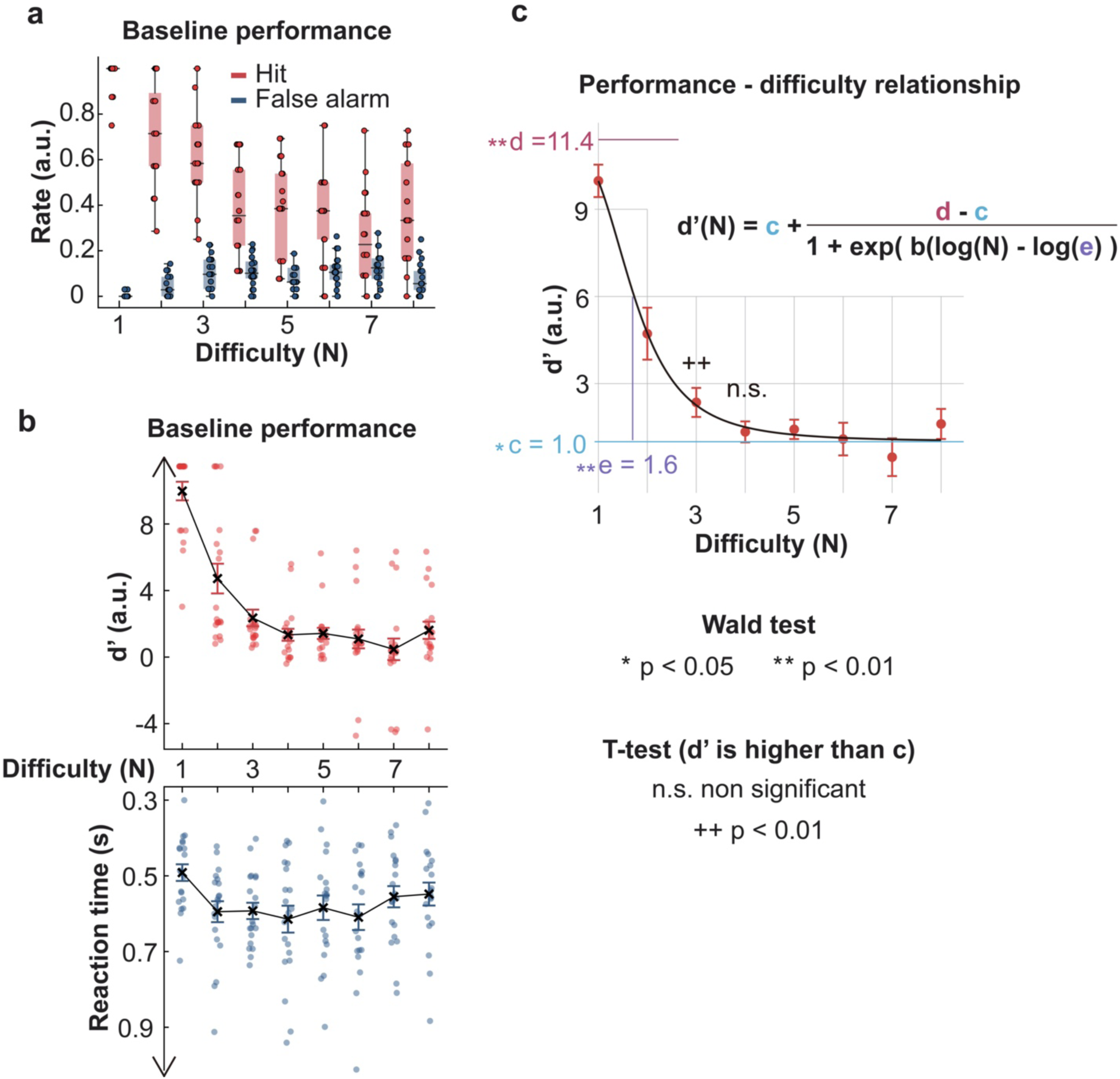
Behavioral outcome at baseline. **a**. The scatters represent the data from 20 subjects. **b**. The mean of *d*′ and reaction time at different difficulties are represented by the cross. The bars represent the standard error. **c**. Nonlinear 4-parameter log-logistic fit of *d*’ as a function of difficulties. The significance of the parameters is indicated with *. T-tests were used to determine if *d*’ is significantly higher than the parameter *c*.

### 3.2 The effects of vibrotactile taVNS on working memory performance

We compared the performance of 4-back tasks to reveal the effect of vibrotactile taVNS on cognitively demanding tasks (**Figure *3*a**). Paired Wilcoxon tests showed that vibrotactile VNS led to a 96% improvement in *d*’ calculated as a percentage change of mean *d*’ compared with the baseline session (Cohen’s d = 0.63, Bonferroni-corrected p-value < 0.01, n = 20). *d*’ during sham sessions was not significantly different from *d*’ during baseline sessions (Cohen’s d = 0.26, Bonferroni-corrected p-value = 0.21). Although *d*’ during VNS sessions is higher than *d*’ during sham sessions, the difference is not significant (Cohen’s d = 0.48, Bonferroni-corrected p-value = 0.78, n = 20). Increased *d*’ at the 4-back task was not associated with increased reaction time (**Figure *3*b**).

We used linear regression to identify the effects of session, session order, and difficulty on *d*’. The model with the lowest AIC was a linear mixed-effect model that incorporates difficulty, session, order, and the interaction between session and difficulty as fixed effects and a random intercept to account for individual differences, shown in equation 3. The median of scaled residuals is 0, and the interquartile range goes from -0.70 to 0.69. T-tests with Satterthwaite’s method showed significant effects of difficulty (95% confidence interval (CI) of *β_N_* = [-3.36, -2.30], p < 0.01), order (95% CI of *β_order_* = 0.29, 1.15], p < 0.01), the interaction between taVNS and difficulty (95% CI of *β_N:VNS_* = [0.19, 1.68], p = 0.01) and taVNS (95% CI of *β_VNS_* = [-4.39, -0.29], p = 0.03) (**Figure *3*c**). Specifically, d’ decreased with increasing difficulty and in earlier sessions (**Supplementary Figure 3**). The significant positive *β_N:VNS_* indicated that the effect of difficulty in *d*’ was attenuated during the VNS session in comparison to the baseline. To test if the attenuation during the VNS session was significant when compared to the sham session, we conducted a one-sided z-test comparing *β_N:VNS_* and *β_N:sham_* . The results showed that *β_N:VNS_* was significantly higher than *β_N:sham_* (z = 1.665, p = 0.048), suggesting that vibrotactile taVNS mitigated the reduction of *d*’ due to increased difficulty when compared both to baseline and sham. The same linear mixed-effect model fitted with d’ data from VNS and sham sessions is shown in **Supplementary Figure 3a**. Besides, the negative *β_VNS_* indicates that vibrotactile VNS might not be helpful for tasks where maximum working memory is not required. For reaction time, the model with the lowest AIC included difficulty, session, and session order as fixed effects and a random intercept to account for individual differences (**Supplementary Table 2**). Our analysis revealed that only difficulty and order significantly influenced reaction time which increased with difficulty and decreased as the session order progressed (**Figure *3*d** and **Supplementary Figure 3**).

**Figure 3.**
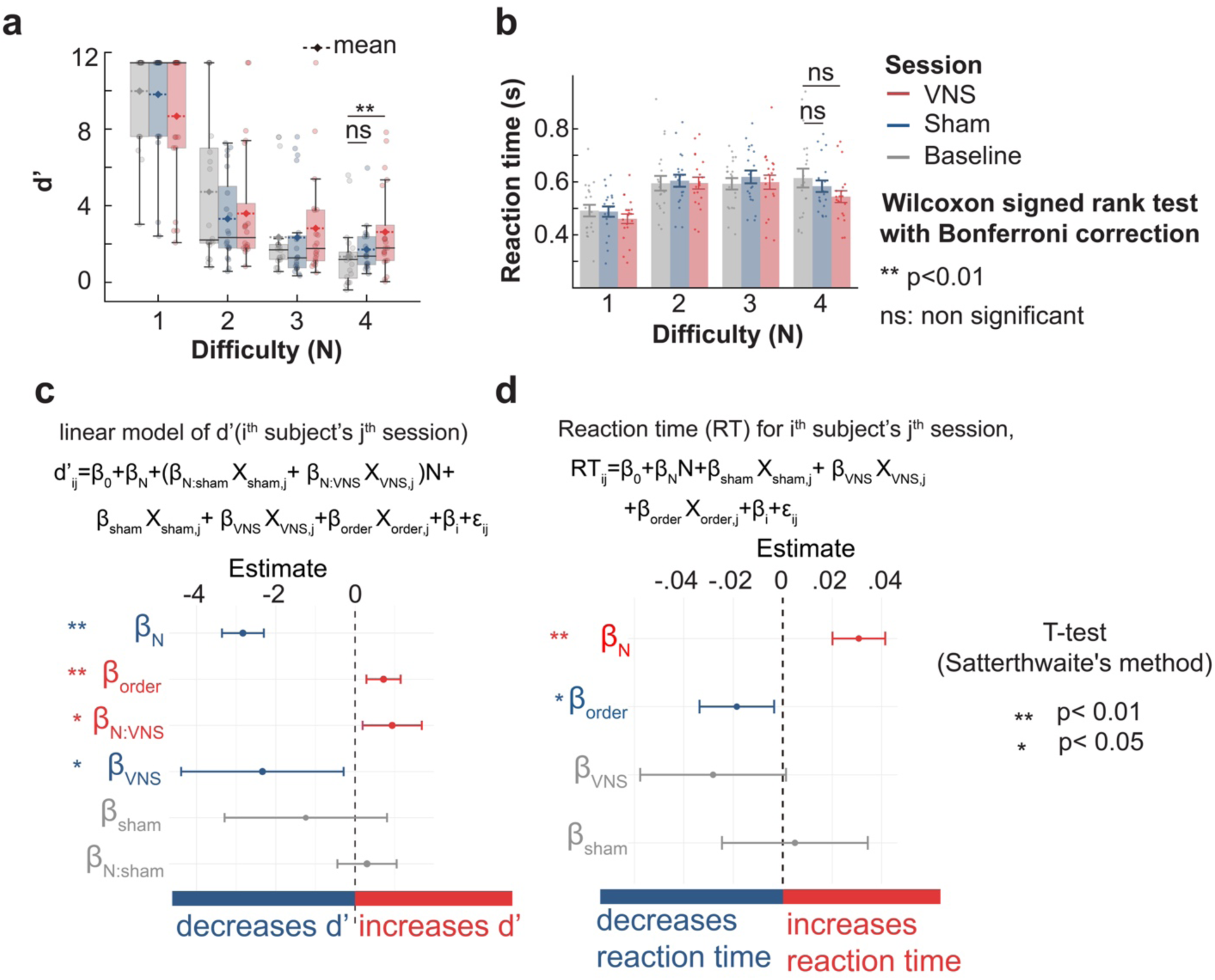
The effects of vibrotactile taVNS on working memory performance. **a, b**. *d*’ and reaction time between sessions at different difficulty levels. The Bonferroni-corrected p-value is 0.005 for *d*’ between VNS and baseline (Test statistics V=182, n=20) and 0.13 for *d*’ between sham and baseline (Test statistics V=154, n=20). Ns indicates that the p-value is larger than 0.05. **c, d**. The estimated coefficients’ values were derived from a linear mixed-effects model of *d*’ and reaction time with the lowest AIC. Positive/Negative estimated coefficients shown in red/blue indicate that *d*’ or reaction time increases/decreases with the corresponding variables. T-tests using Satterthwaite’s method were used to compute the significance of the effects.

### 3.3 Vibrotactile taVNS reduces general arousal decrease due to higher cognitive load or prolonged cognitive demand

We observed that skin conductance increased before the N-back task, which might represent mental preparation for cognitive demand. (**Figure *4*a**). To study the effect of vibrotactile VNS on the temporal dynamic of skin conductance, we rescaled skin conductance data for each N-back from 0 to 1 and aligned them at task onset. We found that the reduction of skin conductance was less steep during the VNS session, indicating that vibrotactile VNS rescued the withdrawal of engagement due to prolonged cognitive demand (**Figure *4*b**). We found that the difference in normalized skin conductance between VNS and sham sessions was significant at the middle and at the end of the N-back task. Non-trivial effect size (Cohen’s d > 0.2) was found 15 seconds after the task onset (**Figure *4*c**). The normalized skin conductance during the rest period between N-back tasks for the three sessions was similar (see **Supplementary Figure 4**). Further, we compared normalized skin conductance between sessions for the three time intervals representing the early, middle, and late stages of the task, respectively (**Figure *4*d**). The normalized skin conductance during VNS was higher than baseline and sham sessions during the middle (Cohen’s d (VNS-sham) = 0.24; Cohen’s d (VNS-baseline) = 0.00) and late stage (Cohen’s d (VNS-sham) = 0.30; Cohen’s d (VNS-baseline) = 0.16).

To study the effect of vibrotactile VNS on general arousal and task engagement in tasks with various difficulties, we normalized the skin conductance level sampled every 10 seconds to the 1-back task in each session to enable comparison across subjects and sessions. The normalized skin conductance level was higher in VNS sessions when the maximum working memory span was reached (**Figure *4*e**). Specifically, in the 3-back task, the normalized skin conductance level in the VNS session was higher compared to baseline but not sham (permutation test, p = 0.02, Cohen’s d = 0.22, Bonferroni-corrected p = 0.08). Normalized skin conductance level was significantly higher in VNS in the 4-back task (permutation test, Bonferroni-corrected p = 0.03 and Cohen’s d = 0.23 for the comparison between VNS and sham session; Bonferroni-corrected p = 0.02 and Cohen’s d = 0.27 for the comparison between VNS and baseline session). To test if the general arousal contributed to the task performance, we grouped baseline-corrected performance metric *d*’ based on the skin conductance level in N-back tasks. For each N-back task of the same difficulty, the mean normalized skin conductance during baseline was subtracted from the mean normalized skin conductance during VNS and sham sessions. We conducted the same procedure on *d*’. We found an optimal skin conductance level for working memory task performance (**Figure *4*f**). Baseline-corrected *d*’ was significantly higher when the baseline-corrected skin conductance level was located above the mean skin conductance level but below one standard deviation of the skin conductance level in the baseline. Cohen’s d of baseline-corrected *d*’ between [−*σ*, 0] and [0, *σ*] baseline-corrected skin conductance level is -0.49, and Bonferroni-corrected p is 0.02. Besides, the effect size of baseline-corrected *d*’ between [0, *α*] and [*α*, 2*α*] baseline-corrected skin conductance level is 0.91, and Bonferroni-corrected p is 0.02. We also found that when the baseline-corrected skin conductance level was within [0, *σ*], the mean hit rate was highest, the mean false alarm rate was lowest, while the mean reaction time was lowest, compared to other skin conductance levels (**Supplementary Figure 4**).

**Figure 4.**
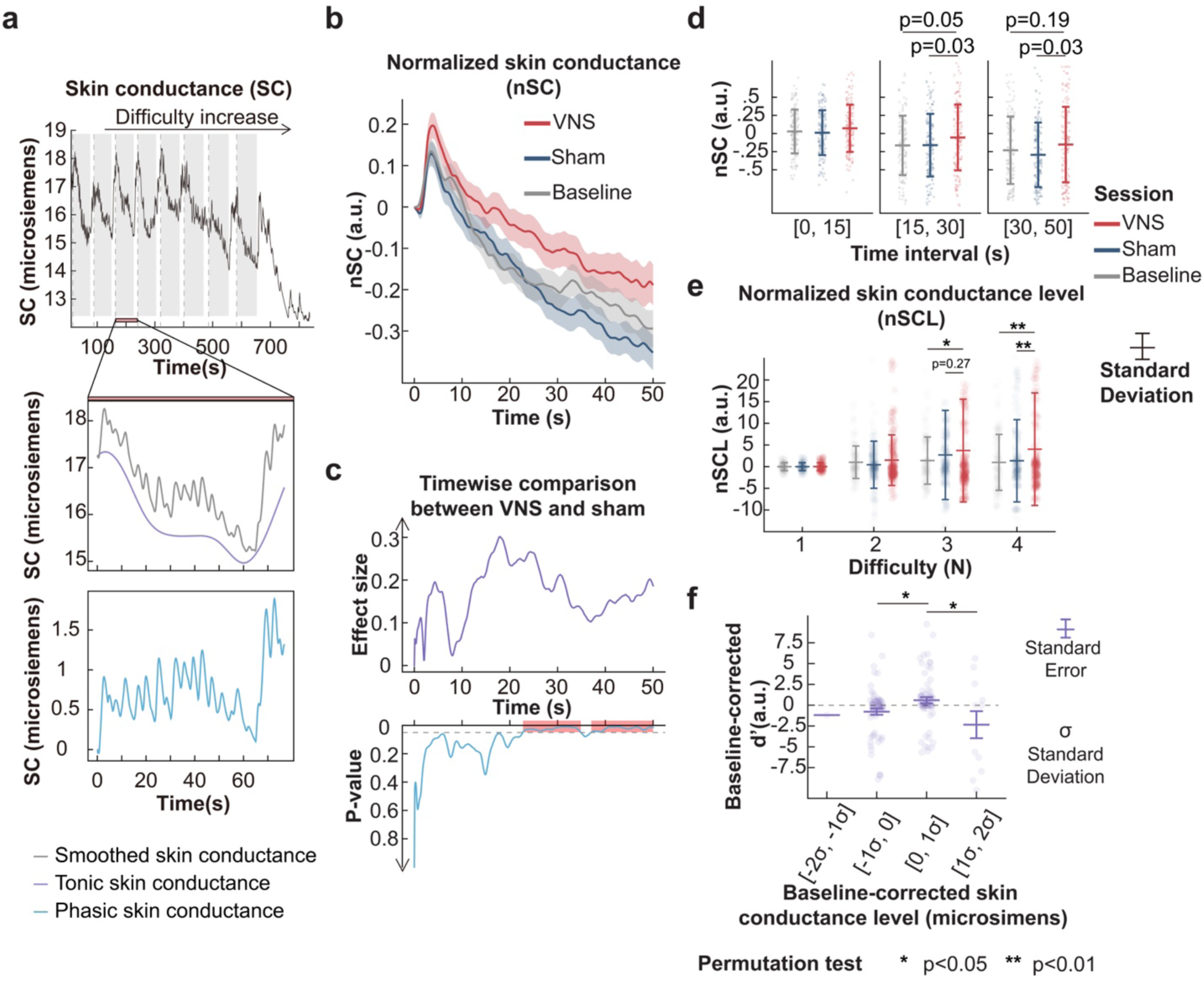
General arousal is modulated by vibrotactile taVNS. **a.** Skin conductance recording for a representative subject during baseline. The vertical dashed lines represent the onset of N-back tasks. The shades represent the task period. The raw skin conductance data during each N-bask task was smoothed and decomposed into a tonic component (i.e., skin conductance level) and a phasic component. The decomposition of the 3-back task is shown. **b.** The normalized skin conductance during the task period over time for the three conditions. **c.** The effect size (Cohen’s d) and p-value of timewise t-test comparison of normalized skin conductance between vibrotactile taVNS and sham sessions. **d.** Normalized smoothed skin conductance under the three conditions for three time intervals identified by timewise comparison. **e.** Normalized skin conductance level (i.e., tonic component) for different sessions during N-back. The permutation test is used to derive p-values in **d** and **e**. Three horizontal bars from top to bottom for each column represent mean + standard deviation, mean, and mean – standard deviation, respectively. **f.** Baseline-corrected d’ from sham and VNS sessions were regrouped based on baseline-corrected skin conductance level distribution.

### 3.4 Evidence from eye-tracking supports that vibrotactile taVNS counteracts decreased arousal during working memory tasks

The left and right pupils changed similarly (Pearson correlation coefficient is 0.93 with a 0.04 standard deviation, see **Supplementary Figure 5**). We used the right pupil diameter for subsequent analyses. The preprocessed pupil diameter showed two components: a 1.5 Hz oscillatory component representing the effect of the stimulus size and luminance of or the transient effect of vibrotactile taVNS and a general trend representing the general arousal (***Figure 5*a**).

The general trend in VNS sessions was higher than in other sessions (***Figure 5*b**). We used a 0.1 Hz low-pass filter to extract the general arousal change. The arousal level rose to a higher level after the task began and decreased less steeply during the VNS session (***Figure 5*c**). A timewise comparison between VNS and sham sessions with t-tests showed that the pupil diameter was higher in VNS after the task began. The difference reached the first peak 10 s after the task onset (***Figure 5*c**). The temporal dynamic of pupil diameter was further broken down into three time intervals. Permutation tests showed that normalized pupil diameter during VNS was higher than other sessions in the [1 s, 10 s] interval (between VNS and Baseline session: p-value = 0.01, Cohen’s d=0.35; between VNS and Sham: p-value = 0.04, Cohen’s d = 0.29, ***Figure 5*d**). To reveal the effect of vibrotactile taVNS during tasks with different difficulties, we subtracted the mean of the filtered pupil diameter during the 1-back task from the filtered pupil diameter for each session. Consistent with the findings in skin conductance, the baseline-corrected pupil diameter was higher in the VNS session in the 4-back task (***Figure 5*e**, permutation test, Bonferroni-corrected p < 0.01 and Cohen’s d = 0.25 for the comparison between VNS and sham session; Bonferroni-corrected p < 0.01 and Cohen’s d = 0.32 for the comparison between VNS and baseline session).

**Figure 5.**
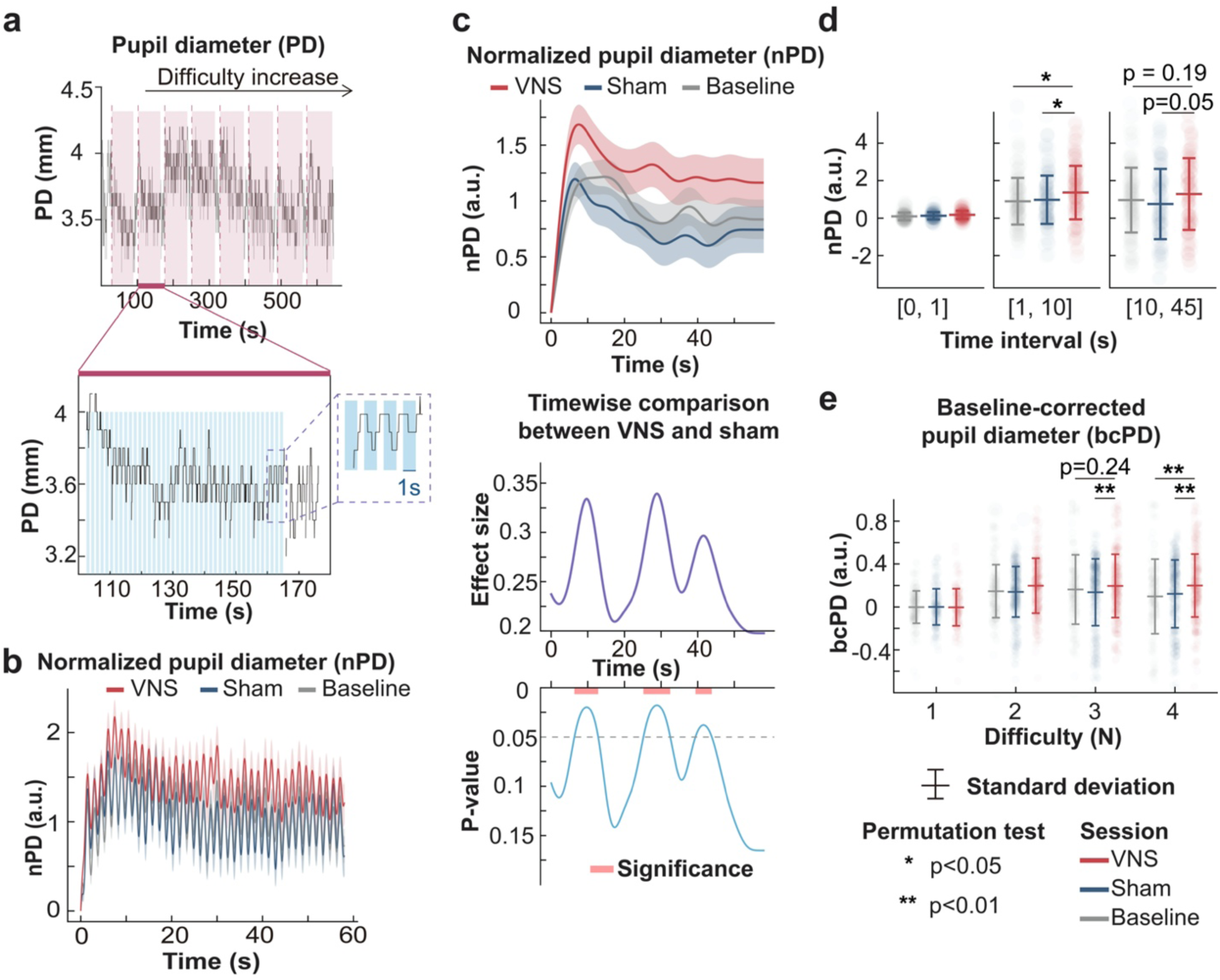
General arousal modulated by vibrotactile taVNS is supported by pupil diameter analysis. **a**. Pupil diameter recording for a representative subject during baseline. The vertical dashed lines represent the onset of N-back tasks. The shades represent the task period. The raw pupil diameter during 2-bask is zoomed in to show that pupil diameter decreases after digit presentation and rebounds after fixation cross representation, indicating that the digit’s luminance and size affect pupil diameter. The blue area represents the time when digits were presented on the screen. A proportion of the pupil diameter is further enlarged for illustration. **b.** The z-score-normalized pupil diameter after 1 Hz low-pass filtering. **c.** The z-score-normalized pupil diameter after 0.1 Hz low-pass filter is used to quantify the general arousal. **d.** Normalized pupil diameter under the three conditions during three time intervals. **e.** Pupil diameter is corrected by subtracting the mean pupil diameter during the 1-back task. We used the permutation test to derive p-values in **d** and **e**. Three horizontal bars from top to bottom for each column represent mean + standard deviation, mean, and mean – standard deviation.

## Discussion

In this initial investigation, we examined the capacity of vibrotactile vagus nerve stimulation to augment working memory. Our evidence suggests that applying vibrotactile taVNS is most efficacious when the cognitive load approaches the individual’s limit. The mechanism appears to involve stabilizing arousal levels that typically wane during sustained working memory tasks. Moreover, vibrotactile taVNS seemed to elevate and maintain arousal levels to a state conducive to working memory function, making it potentially valuable for conditions where working memory is compromised by reduced arousal. A study demonstrated that taVNS improves human working memory under sleep deprivation stress^11^. Our study also illuminates the promise of vibrotactile taVNS as an intervention for patients with neurologically impaired working memory due to diagnoses such as Alzheimer’s disease or brain injury. Targeted research in patient populations will be necessary to ascertain its therapeutic benefits.

Insight into the functional and physiologic implications of vibrotactile taVNS is in its infancy. Pupil diameter has been shown to track noradrenergic locus coeruleus and cholinergic basal forebrain activity^29, 30^. Our findings suggest that vibrotactile taVNS mimics the effect of invasive VNS and activates the locus coeruleus-noradrenaline system. In addition, VNS is thought to increase cortical inhibition through the release of gamma-aminobutyric acid (GABA) from the nucleus of the solitary tract^31^. This process is believed to sharpen task-relevant representations and improve working memory performance^32,33^. Neuroimaging studies showed that deactivation of the ventromedial prefrontal cortex was associated with elevated skin conductance levels^34^. In our study, skin conductance level was higher when subjects received active vibrotactile taVNS, suggesting vibrotactile taVNS may also lead to ventromedial prefrontal cortex deactivation.

A major limitation of this study is that the results of the behavioral effects of vibrotactile taVNS are inconclusive. Our findings support that performance during active stimulation is significantly higher than baseline control in the 4-back task. However, the direct comparison of *d*’ between active stimulation and sham control did not yield significant results. These results allow for four potential hypotheses: (1) Stimulating the concha does not differ significantly from earlobe stimulation. (2) The auricular branch of the vagus nerve was activated during sham-control condition as mechanical stimulation may be more widespread than classically applied bipolar currents in electrical stimulation. (3) The study may lack sufficient statistical power to discern differences between active stimulation and sham control. In our study, the effect size between active stimulation and sham control is 0.26. Therefore, 124 subjects are needed for paired Wilcoxon signed-rank test to reach 80% power, assuming the type I error rate is 0.05. (4) The imbalance in the number of sham sessions (nine) versus VNS sessions (six) that are the third session potentially confounds outcomes. To conclusively determine whether vibrotactile taVNS facilitates working memory, replicating this pilot study with a more substantial sample size is needed. The application of vibrotactile taVNS will also benefit from investigating vibrotactile taVNS of systematically varying frequency.

## Code availability

https://github.com/neurotechcenter/taVNS_physiology

## CRediT authorship contribution statement

**Gansheng Tan**: Conceptualization, Methodology, Formal analysis, Data curation, Writing – original draft, Writing – review & editing. **Josh Adams**: Stimulation device, Writing – review & editing. **Kara Donovan**: Methodology, Writing – review & editing. **Phillip Demarest**: Methodology, Writing – review & editing. **Jon T. Willie**: Methodology, Writing – review & editing. **Peter Brunner**: Supervision, Experimental platform setup, Methodology, Writing – review & editing. **Jenna L. Gorlewicz**: Stimulation device, Writing – review & editing. **Eric C. Leuthardt**: Conceptualization, Supervision, Methodology, Writing – review & editing, Writing – original draft.

## Declaration of generative AI and AI-assisted technologies in the writing process

While preparing this work, the authors used chatGPT to check grammar errors. After using this tool, the authors reviewed and edited the content as needed and took full responsibility for the content of the publication.

## Acknowledgments

We thank the subjects who volunteered to participate in the study. We acknowledge Kay Park and Sid Sivakumar for their role in subject recruitment. We thank Drs. Hohyun Cho and Tao Xie for sharing their expertise. We thank Dr. Paul Cassidy for his contributions to the scientific editing of this manuscript, supported by the Institute of Clinical and Translational Sciences grant UL1TR002345 from the National Center for Advancing Translational Sciences (NCATS).

