## Supplementary Figure 1 for "Does Vibrotactile Stimulation of the Auricular Vagus Nerve Enhance Working Memory? A Behavioral and Physiological Investigation"

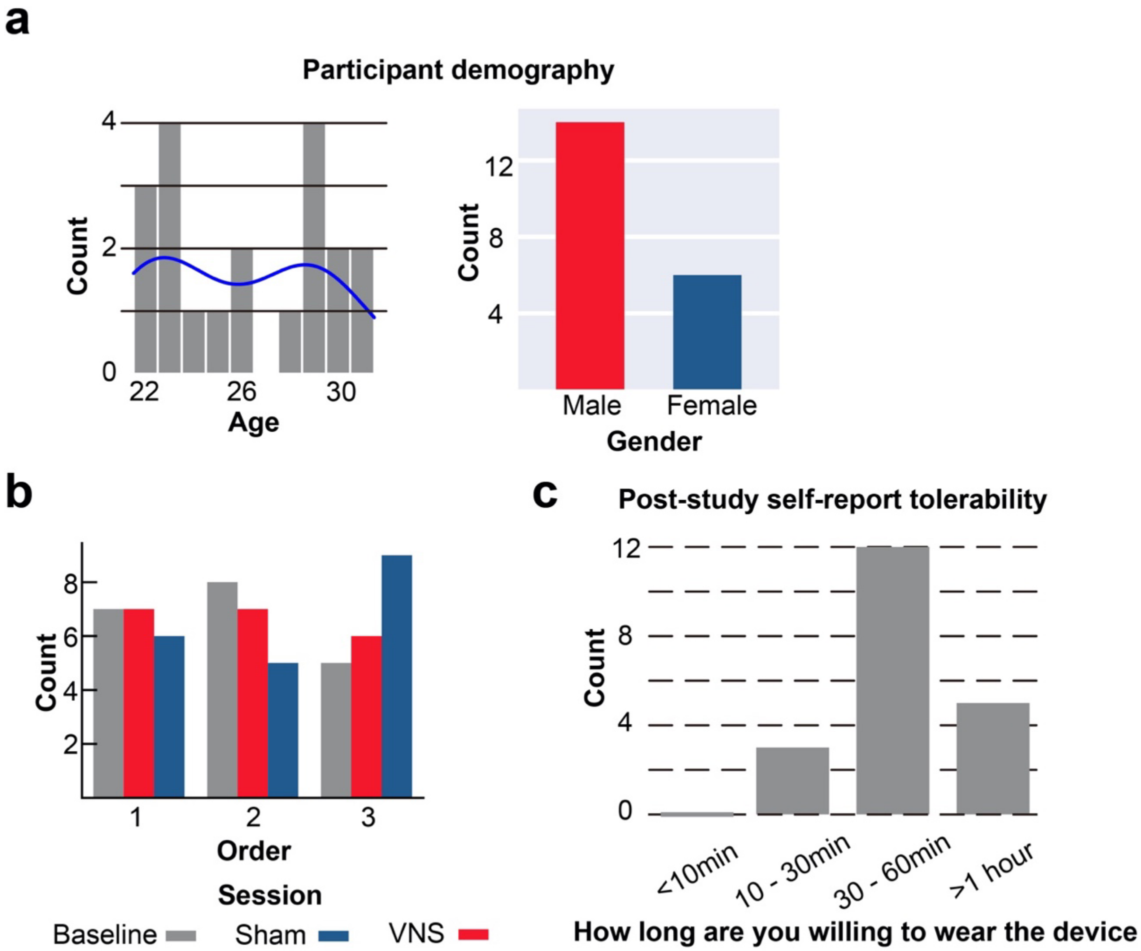


**Supplementary Figure 1** (related to Figure 1) Subject demographics, session order summary, and subject feedback about the vibrotactile taVNS device. **a**. The histogram of age and gender for recruited subjects. **b**. The y-axis represents the number of baseline, sham, and VNS sessions at different orders. The session order was randomized by generating 20 random numbers between 1 and 6. Each number represents a specific baseline, sham, and VNS session order. **c**. the histogram about the answer to the question: How long are you willing to wear the vibrotactile taVNS device? Out of the 22 participants initially recruited, two were excluded. One subject was excluded from the analysis because the subject developed hard skin from rock climbing, preventing skin conductance recording. Another subject was excluded from the analysis because the subject reported that the vibrotactile taVNS did not contact the concha region during one of the sessions. The sample size was calculated to achieve 80% power based on the effect size reported in a comparable study that examined the impact of taVNS on hit rates in 3-back tasks[1]. The aforementioned study showed the following hit rate statistics: during the online taVNS – baseline session, the mean hit rate was 0.757 with a standard deviation (SD) of 0.021; during the online taVNS – baseline session, the mean hit rate was 0.766 with an SD of 0.021. During the sham – baseline session, the mean hit rate was 0.758 with an SD of 0.021, and during the sham – stimulation session, the mean hit rate was 0.751 with an SD of 0.026. The mean difference in hit rates between the stimulation session and the baseline session was 0.009, and the standard deviation was 0.021 for the online taVNS condition. The mean difference in hit rates was -0.007, and the standard deviation was assumed to be 0.026 for the sham condition. Cohen's d is calculated as $\frac{0.009-(-0.007)}{\text{pooled standard deviation}} = \frac{0.009-(-0.007)}{\sqrt{0.021^2+0.026^2}} = 0.677$. With an anticipated effect size of 0.677 and a Type I error rate (α) of 0.05, a paired two-tailed t-test with a sample size of 20 is estimated to detect the difference between the VNS and sham conditions with 82% power. Of the 20 participants included in the analysis, 10 began the task between 10:30 a.m. and 12:30 a.m., while the other 10 began the task between 1:30 p.m. and 5:30 p.m.. Due to logistical constraints, our study shared a limited number of eye trackers with another ongoing research project focused on epilepsy patients. Unforeseen scheduling conflicts with the epilepsy research project arose. These conflicts resulted in the unavailability of eye trackers for 3 of our subjects at their scheduled experiment times. Consequently, eye tracking data were successfully collected for 17 of the recruited subjects.

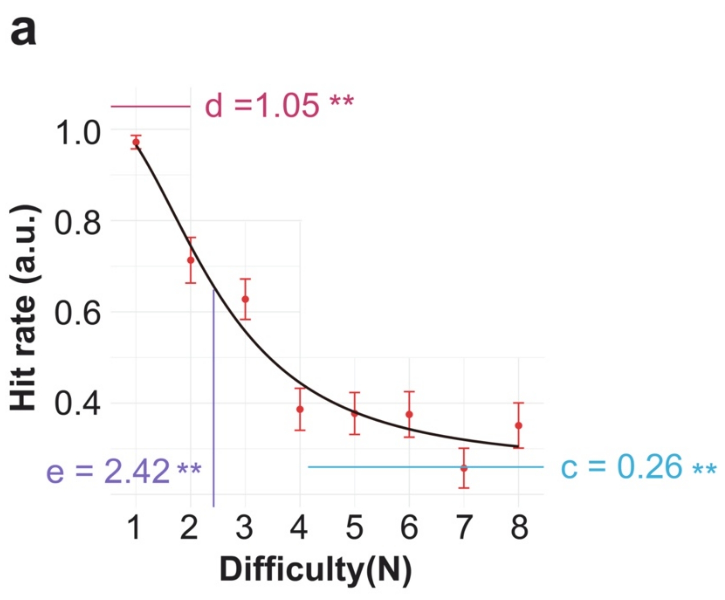


**Supplementary Figure 2** (related to Figure 2) **a**. The 4-parameter log-logistic regression model shows that the maximum working memory span is most likely to be reached at N=4. **b**. The proportion of positive probes for each difficulty.

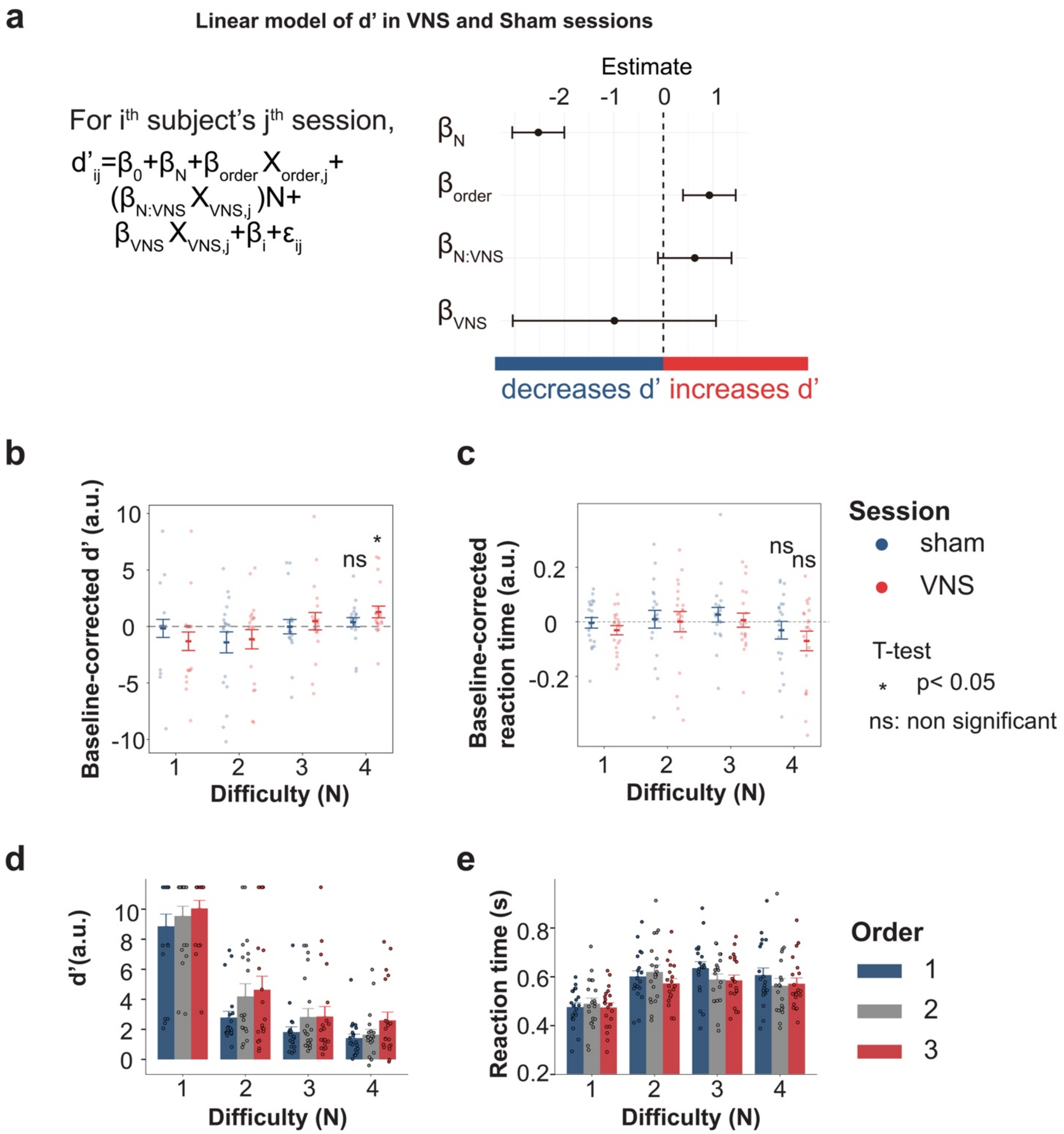


**Supplementary Figure 3** (related to Figure 3) **a**. The linear mixed-effect model fitted with $d'$ data from VNS and sham sessions. 95% CI of $\beta_N$ is [-3.05, -2.00]. 95% CI of $\beta_{order}$ is [0.40, 1.46]. 95% CI of $\beta_{N:VNS}$ $is$ [-0.11, 1.38]. A one-sided t-test using Satterthwaite approximation of the degree of freedom showed that $\beta_{N:VNS}$ was significantly larger than 0 (p-value = 0.048). 95% CI of $\beta_{VNS}$ is [-3.04, 1.07]. **b**. Baseline-corrected $d'$ at N=4 is significantly higher than 0 during the VNS session. **C**. Reaction time in VNS and sham was not significantly different than baseline. **d-e**. The effect of session order on $d'$ and reaction time.

| | Formula | AIC |
|---|---|---|
| Model 1 | $d' \sim n * session + order + (1\| subject)$ | 1198 |
| Model 2 | $d' \sim n + session + order + (1\| subject)$ | 1200 |
| Model 3 | $d' \sim n * session + order$ | 1213 |
| Model 4 | $d' \sim n + session + order$ | 1214 |
| Model 4 | $d' \sim n + session$ | 1220 |

**Supplementary Table 1** Model selection for $d'$.

| | Formula | AIC |
|---|---|---|
| Model 1 | $reaction\ time \sim n + session + order + (1\| subject)$ | -404 |
| Model 2 | $reaction\ time \sim n + order + (1\| subject)$ | -403 |
| Model 3 | $reaction\ time \sim n * session + order + (1\| subject)$ | -401 |
| Model 4 | $reaction\ time \sim n + session + order$ | -364 |
| Model 4 | $reaction\ time \sim n + order$ | -364 |

**Supplementary Table 2** Model selection for $reaction\ time$.

| Settings | Color | Size/orientation |
|---|---|---|
| Background screen | Black (#231F20) | |
| Fixation cross | Red (#ED1C24) | The fixation cross was composed of a horizontal and a vertical arm, each measuring 90 points in width and 8 points in height assuming the monitor is of 2560*1440 points. |

| | | |
|---|---|---|
| Digits | White (#FFFFFF) | The digit size is relative to the monitor dimension. In a monitor of 2560*1440 points, the digits were rendered in Arial font at a size of 300 points. |

**Supplementary Table 3** Stimuli information. The experiment was conducted in standard office ambient lighting conditions (150-600 lux).

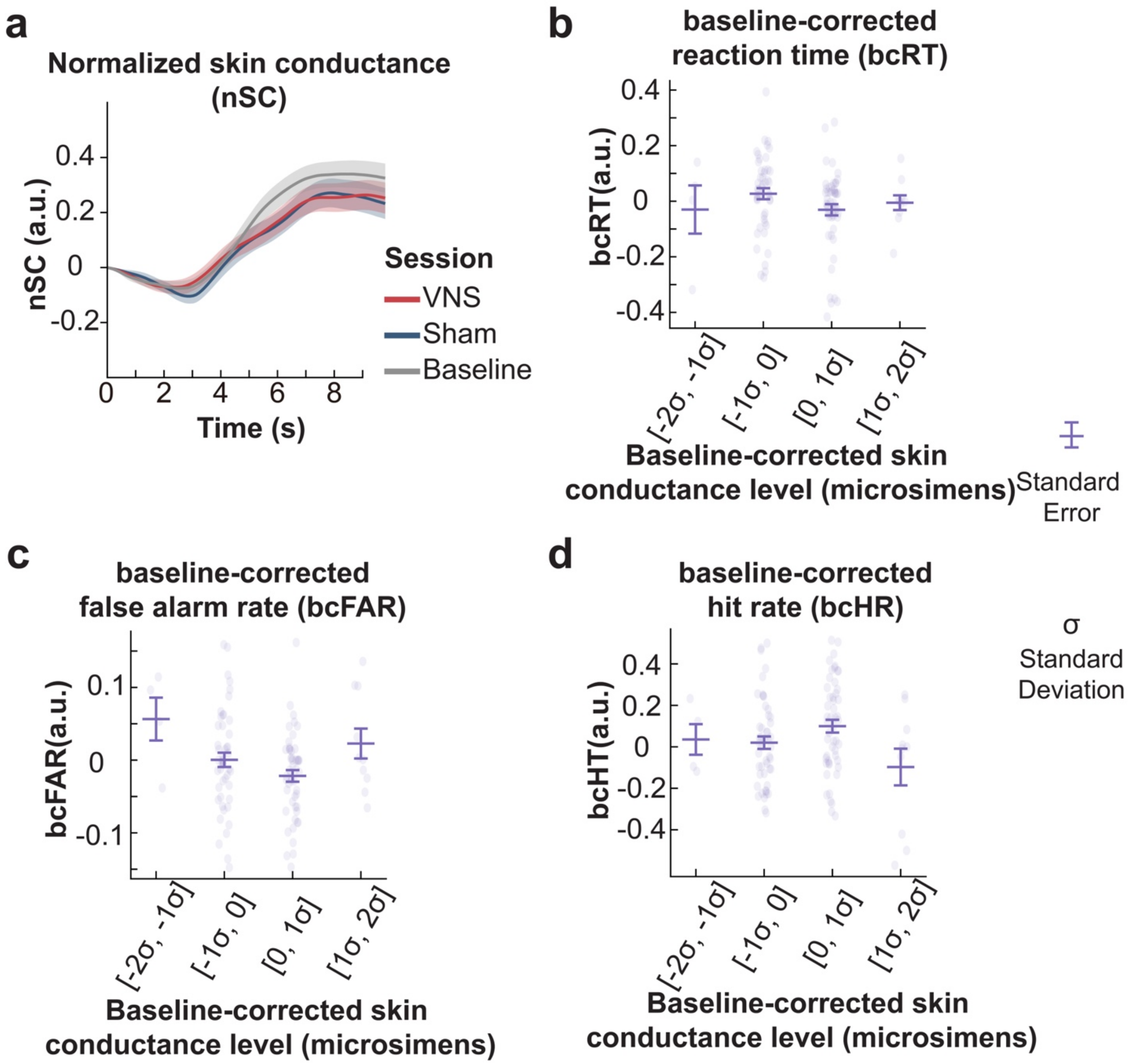


**Supplementary Figure 4** (related to Figure 4) **a**. Normalized skin conductance during the rest intervals between two N-back tasks. Time 0 represents the onset of rest between tasks. **b**. The relation between baseline-corrected reaction time and baseline-corrected skin conductance level in sham and VNS sessions. **c**. The relation between baseline-corrected false alarm rate and baseline-corrected skin conductance level in sham and VNS sessions. **d**. The relation between baseline-corrected hit rate and baseline-corrected skin conductance level in sham and VNS sessions.

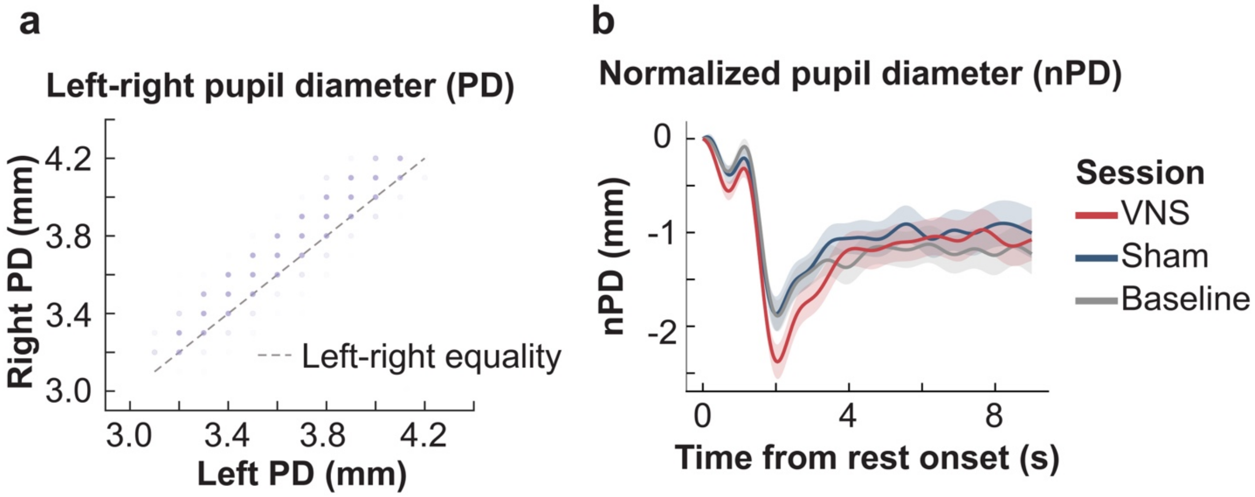


**Supplementary Figure 5** (related to Figure 5) **a**. Right pupil diameter against left pupil diameter for the representative subject in Figure 5. Each point represents the pair of right pupil diameter and left pupil diameter at a given time point. The mean absolute deviation is 0.19 with a 0.11 standard deviation, and the mean Pearson correlation coefficient is 0.93 with a 0.04 standard deviation. **b**. Normalized pupil diameter during the rest periods following each N-back task. After completing a N-back task, subjects were allowed to initiate the subsequent N-back task at their pace, with a mandatory minimum rest interval of 10 seconds. Time 0 represents the onset of the rest intervals between N-back tasks.
